# Decrease in reaction time for volleyball athletes during saccadic eye movement task: An associative analysis with evoked potentials

**DOI:** 10.1101/2023.08.03.551793

**Authors:** Élida Costa, Mariana Gongora, Juliana Bittencourt, Victor Marinho, Mauricio Cagy, Silmar Teixeira, Eduardo Nicoliche, Isabelle Fernandes, Jacob Wienecke, Pedro Ribeiro, Daya S. Gupta, Bruna Velasques, Henning Budde

## Abstract

**Aim:** The current study investigated the differences to event-related potential and reaction time under two groups (athletes *vs*. non-athletes).

**Material and Methods:** The P300 was analyzed for Fz, Cz and Pz electrodes in thirty-one healthy volunteers divided into two groups (volleyball athletes and non-athletes). In addition, the participants performed a saccadic eye movement task to measure reaction time.

**Results:** The EEG analysis showed that the athletes in comparison to the no-athletes has differences of the P300 in the frontal area (p=0.021). In relation to reaction time, the results show lower reaction time for athletes (p=0.001).

**Conclusions:** The volleyball athletes may present a greater allocation of attention during the execution of the inhibition task, since they have a lower reaction time for responses when compared to non-athletes.

## 1. Introduction

Skilled performance is a key role in improving cognitive aspects such as attention, memory and decision-making [1,2]. According to a previous study, it was possible to identify neural correlates in volleyball athletes, the findings demonstrated that volleyball training is related to lower latency of evoked potentials [3]. Some neuroimaging and neurophysiological studies in athletes have shown that neural processing during cognition and decision-making for motor act are modulated by long-term perceptual motor training and may present patterns of neural inputs and outputs consistent with neural efficiency [4,5,6,7]. Although it is, evident those sports are related to physical and mental benefits, it is necessary to understand how regular sports practice interfere improves cognitive processes [8,9]. Thus, can provide a more comprehensive understanding of the differences in relation to neural processing for attentional level, with greater focus on high-performance athletes when compared to non-athletes and beginning athletes [4].

The EEG-analysis by event-related potential (ERP) P300 simultaneously with cognitive tasks are methods to evaluate patterns of cognitive processing, component elicited in the process of decision-making [10,11,12]. The concomitant assessment enhances the functional diagnosis of the central visual pathway, as well as tool for studying the endogenous potential in the cortical mechanisms of visual perceptual processing, since your occurrence links not to the physical attributes of a stimulus, but to a person’s reaction to it [13]. In particular, the P300 supports the evidence about the level and orientation of attention, contextual updating, modulation and response resolution by measures of latency and amplitude [14,15]. The P300 amplitude is directly proportional to the subject’s attention and the increase in the P300 latency may indicate the prolonged temporal processing during complex cognitive information [16,17,18].

Athletes’ cognitive engagement is necessary for high performance in several sports, in focus the volleyball, has been studied extensively [19,20]. The practice of this sport requires a high attention process due to vast exposure to dynamic visual stimuli. Therefore, the ability to have efficient attention is an essential factor for the success of these elite athletes [21]. In the sense, the premotor theory of attention defines the attention processes in response to visual-motor stimuli, which states shifts of attention occur by planning goal-directed actions such as eye movements and reaches. The spatial attention and motor preparation could be structurally and functionally equivalents, and may share neural networks when the situation involves directly planning an action mainly to the oculomotor system [4,14,15,22,23].

Previous studies demonstrated that P300 ERPs components in volleyball athletes were significantly improve when to compare with non-athletes [21,23]. The understanding of this athletes performance during the sport can be supported on the premotor theory assumes that goal-driven by attention level is a dynamic mechanism, where an ocular motor system formed can brings the target into the fovea [24]. The substantial difference between driven “movement” and eye movement can be efficiently measured through saccadic eye movement (SEM) paradigm, because it allows investigating the first stages of visual processing and its relationship with attention [25,26,27]. However, the question of what determines the factors contributing to this relationship remains unanswered.

We hypothesized that athletes would present greater P300 amplitude and shorter reaction time when compared with non-athletes. The state of the art demonstrated that no study has examined the combinatorial relationship of P300 and saccadic go/no-go task for comparisons between volleyball players’ and non-athletes. Thus, the study contributes to knowledge about neural mechanisms underlying attentional processing during sports performance, and may influence the construction of intervention strategies to sports performance and reassert the use of sport as a resource to improve attention performance in non-athletes.

## 2. Material and methods

### 2.1. Participants

We recruited thirty-one healthy volunteers (4 men, 11 women), with age from 12 to 17 years (mean ± standard deviation [SD] = 16.9 ± 0.3 years). The subjects were divided into two groups: sixteen volleyball athletes (15.8 ± 0.2 years) and fifteen non-athletes as a control group (16.2 ± 0.3 years). The volunteers were recruited from June to August 2009. An Independent *t*-test was performed between the two groups for age and showed no significant difference (p>0.05). Only right-handed individuals were selected based on the Edinburgh Inventory [28].

We analyzed the Sustained Attention Test (SA) to investigate if the functions of concentrated attention, the speed with quality, and support of attention were comparable [29].

In the context, an Independent *t*-test was performed between the two groups for SA test and showed no significant difference (p>0.05), with Cronbach’s alpha (α)= 0.79; p=0.001. The results demonstrated no differences for neuropsychological tests of attention (concentration p=0.70 ± 6.5, total hits p= 0.40 ± 4.0; speed with quality p=0.40 ± 8.1).

SA test description: The tests were corrected using the total number of correct answers, errors, and omissions. During the test, participants were submitted to a sequence of visual stimuli (target and non-target figures); and asked to mark the target stimuli with a risk. The test is performed with a pencil and an answer sheet and consists of 25 rows with 25 stimuli each. The subject must select only one type of stimulus among the possibilities. The participant has 15 seconds to complete each row. At the end of the established time, the applicator gives the command to go to the next one, and, in this way, the participant immediately goes to the following line starts again. On average, the application time is 10 minutes.

All participants had normal vision, in addition, were not using any substance that could influence brain activities (e.g. tobacco, coffee, alcoholic beverages, caffeine-containing foods, or medications) 14h before or during the study period. Routine ophthalmologic examinations confirmed that all participants had normal visual function. The participants underwent a medical evaluation to exclude neurological or motor diseases and contraindications to the experimental procedure. The athletes in the study were recruited from the Brazilian national volleyball team and practiced volleyball for around 5.0 ± 2.8 years. The non-athletes not involved in any regular, sports activity.

Ethics approval was obtained, under number CAAE: 94619218.3.000.5257. The participants signed a Free and Informed Consent Declaration under the ethical standards established in the Helsinki Declaration, 1964.

### 2.2. Experimental procedure and task

The participants were accommodated in a room with brightness control, sound isolation, and electrical grounding. A 120 cm length bar composed of 13 light-emitting diodes (LEDs) was placed 100 cm away from the participant’s eye level (Figure 1). The bar had a central warning (bi-color LED – green and red) and six more LEDs located on each side (6 LEDs located on the left side of fixation and 6 LEDs located on the right side). The distance between the participants’ eyes and the LED bar was standardized at 100 cm. The computer software – SEM Acquisition controls the LED bar determining the presentation of the stimulus and measure the reaction time to perceive the visual stimulus. The reaction time recording is associated in combination with the ocular electrical activity, or electrooculogram (EOG), which captured by the placement of two 9 mm diameter electrodes mounted bipolarly. The electrodes were placed in the outer corner of the left and right eyes that recorded the horizontal eye movements (hEOG).

**Figure 1.**
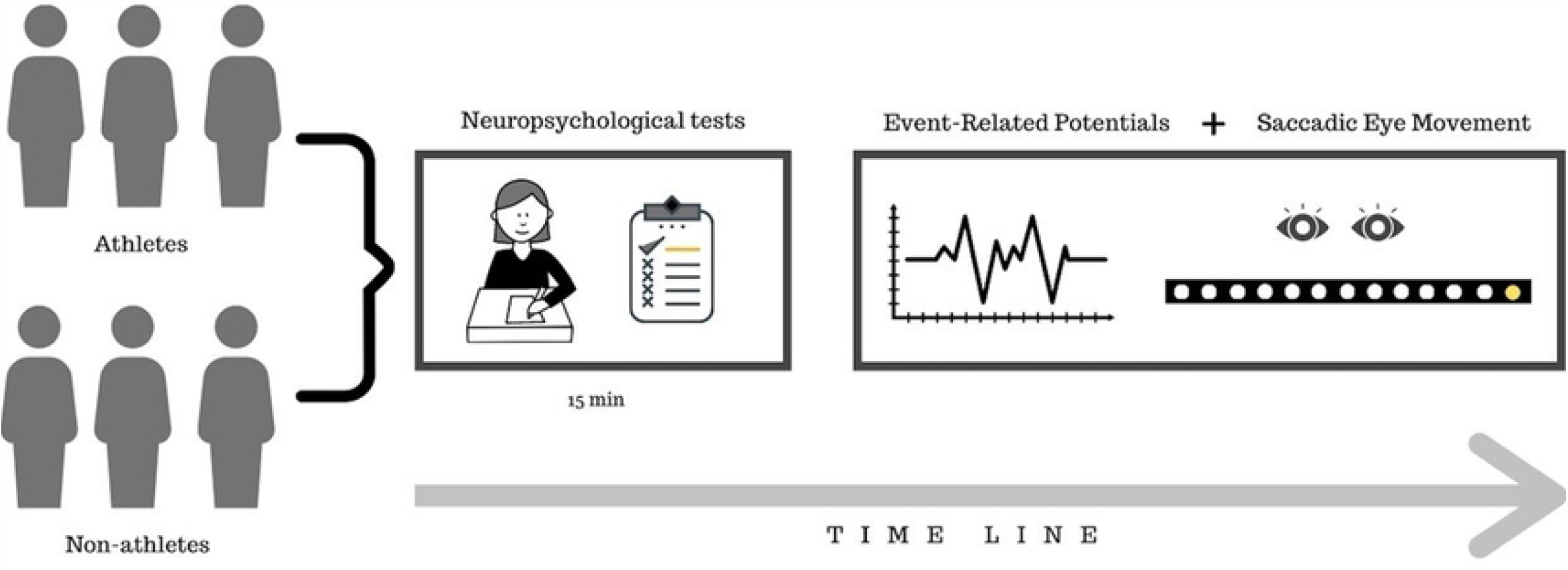
Experimental Design.

We analyze the EEG acquisition during the task execution (Figure 1). All subjects performed one block with 120 trials at their own pace during of the SEM paradigm. The procedure consisted with ten target-LED lights placed on the right side of the bar and 10 LED lights on placed the left side of the bar from the center. During the experimental procedure, the participants were instructed to keep their eyes fixed on the center of the bar and shift their eyes when they perceived one of the diodes lighting up, no head movements, only eye movements. The saccadic eye movement (SEM) paradigm consisted of a fixed pattern of stimulus presentation where the target-stimulus (target-LED) always appeared randomly between left and right sides at a predefined position, this condition was characterized by the predictability of the appearance of the stimulus in time and space, being considered directed by memory. The paradigm is characterized by predictability since the stimulus appears at a predefined spatial location in the periphery of the visual field. Each LED remained lit for 250 ms, with an inter-LED time of 2 seconds.

### 2.3. EEG recording

All subjects were accommodated in a room with acoustic insulation, electrical grounding, and low light. Subjects sat in a chair with armrests to minimize muscle artifact during EEG signal acquisition. The 20-channel continuous EEG was recorded by BrainNet BNT36 (EMSA Medical Equipment). The silver/silver chloride electrodes were positioned through a nylon cap following the international 10–20 system, including binaural reference electrodes (SPES Medical Brazil). The EEG electrodes impedance and EOG electrodes were kept below 5kΩ. The acquired data had an amplitude below 100 μV. The sampling rate was 240 Hz. An antialiasing low-pass filter with a cut-off frequency of 100 Hz was employed. It was configured to use 60 Hz Notch digital filtering, with highpass filters at 0.03 Hz and low pass filters at 40 Hz (Order 2 Butterworth filter), using the Data Acquisition software (Delphi 5.0).

The signal corresponding to each EEG derivation came from the electric potential difference between each electrode and the pre-set reference (earlobes). The epochs were time-locked to the stimulus presentation, and we extracted 15 epochs for each participant before the stimuli appearance and 15 more epochs after the stimuli presentation.

The electrodes located on the central line of the head, Fz, Cz and Pz were selected due to their relationship with the interconnectivity pattern of left and right hemispheres better related to neurobiological process [30,31].

### 2.4. EEG data processing

A visual inspection and independent component analysis (ICA) was applied to identify and remove all remaining artifacts through Matlab 5.3® (The Mathworks, Inc.). Data from individual electrodes that showed contact loss with scalp or high impedance (>5kΩ) were not considered. After ICA, the overall rate of removal for noisy data in each participant was less than ten percentage independent of the task condition. A classical estimator (i.e., parametric, Bartlett Periodogram, using non-overlapping 2 s long [480 samples] rectangular windows) was applied to the Power Spectral Density (PSD), estimated from the Fourier Transform (FT), which was performed using MATLAB (Mathworks, Inc.). Epochs were selected between 1-sec pre-stimulus to 1.5-sec post-stimulus. The total number of epochs used after visual inspection and ICA for each group was as follows: non-athlete group (n= 376 epochs); athlete group (n= 366 epochs).

After specific channels were selected (Fz, Cz, and Pz), the event-related potentials (ERPs) transform was computed for the electrodes. The data were averaged and represented graphically in terms of latency (x-axis) and amplitude (y-axis). In the context, the P300 component was identified as the most positive component within the latency window of 250-500 ms. Amplitude was measured relative to a pre-stimulus baseline, with peak latency defined as the time point of maximum positive amplitude within the specific latency window.

### 2.5. Statistical Analysis

Statistical procedures were conducted using IBM SPSS for Windows (version 21.0; IBM, Armonk, NY, USA). Analyses were controlled for age using an Independent *t*-test, where no significant effects were found. The normality and Homogeneity of variance of the data were previously verified by the Shapiro–Wilk and Levene tests. We use the data as mean, standard deviation (SD) and standard error (SE). The differences in the P300 amplitude and the reaction time for SEM task were analyzed by Independent *t*-tests between athlete and non-athlete groups, with the analysis effect evaluated by the Cohen’s d. The effect sizes were calculated (≤0.039: no effect, 0.04–0.24: minimum, 0.25–0.63: moderate, ≥0.64: strong according to Ferguson (2009). For all statistical analyses, the significance level was α = 0.05 [32,33].

## 3. Results

### 3.1. Reaction Time

The analysis by Independent *t*-test showed statistical difference [*t*(1)= 4.71; p=0.001; d=0.21; CI95%= 1.14 – 2.91], with a lower reaction time in the athlete group (mean: 317.84ms, SD: 58.68ms, SE: 1.92ms) when compared to non-athlete (mean: 330.99ms, SD: 52.49ms, SE: 1.83ms). In addition, the findings revealed a minimum effect for behavioral measure (Figure 2).

**Figure 2.**
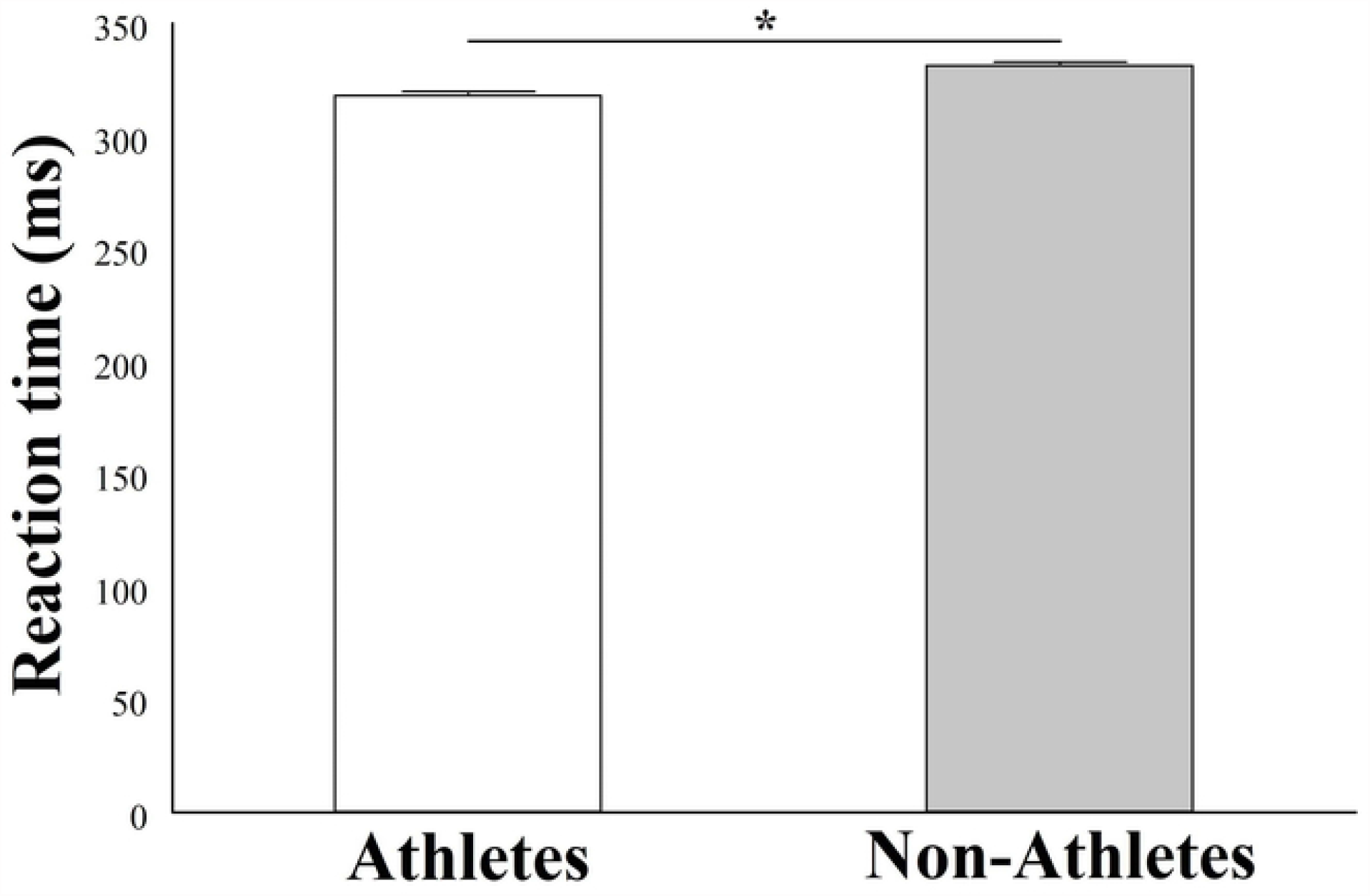
Reaction time for athlete (left) and non-athlete (right) for the saccadic eye movement (SEM) paradigm. Differences of means significant in reaction time were obtained by mean and standard deviation, and the statistically significant differences (p=0.001) are indicated with (*).

### 3.2. P300 event-Related Potentials

The analysis of P300 event-related potentials by Independent t-tests for Fz electrode revealed difference between groups, with [*t*(1)= 4.43; p=0.021; d=0.23; CI95% -0.008 – 0.086] (Figure 3). The atlhetes group had greater mean potential of P300 ERPs when compared to the non-athletes. In relation to Cz and Pz electrodes, no found statistical difference (p>0.05). We demonstrated the target ERPs amplitude of each electrode inspected (Fz, Cz, and Pz). The mean potential (μV) of visual P300 ERPs from the athletes group and non-athletes were demonstrated in response to the visual stimulus (13 light-emitting diodes) for the SEM task (Figure 4, 5 and 6).

**Figure 3.**
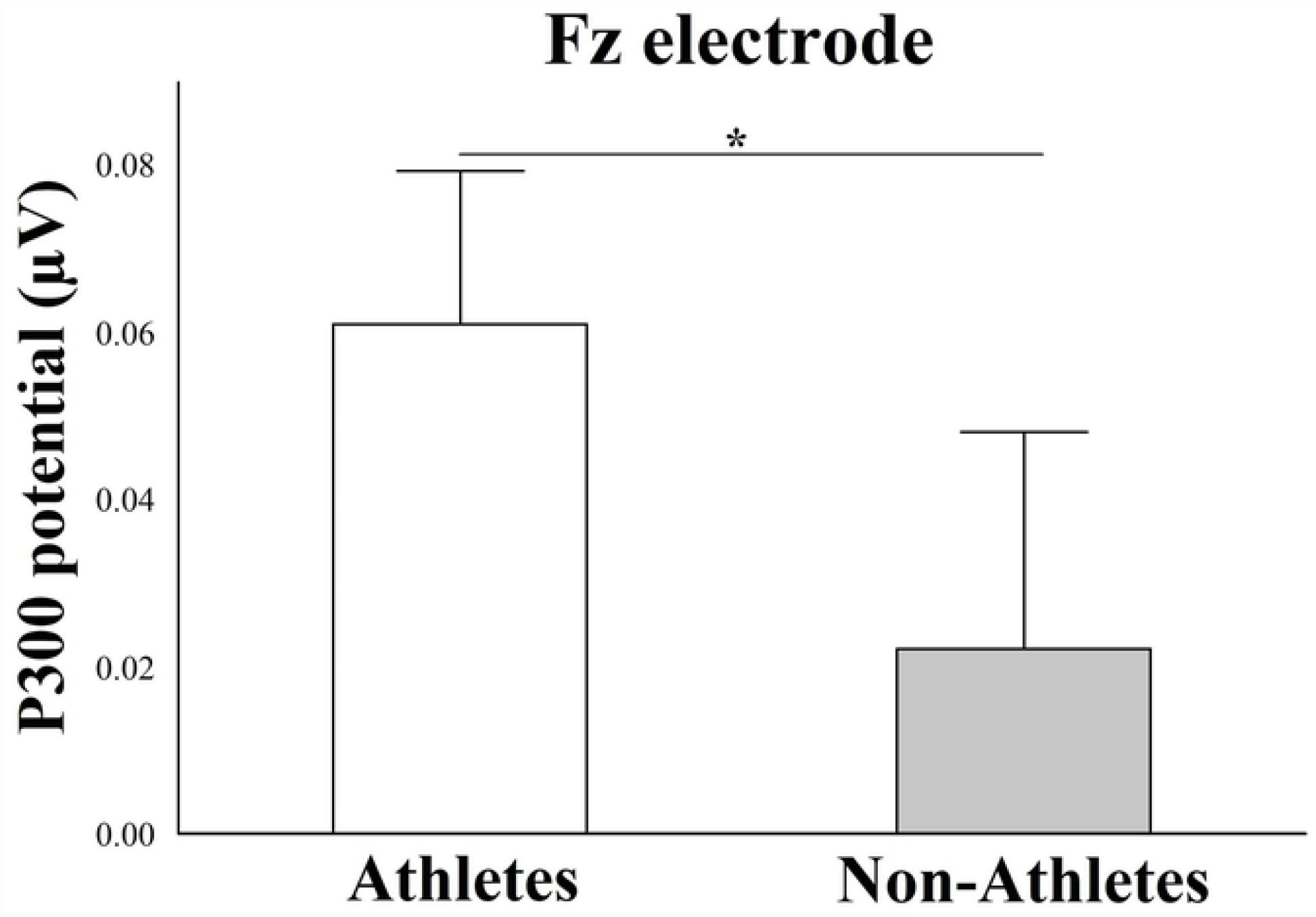
Difference for P300 under Fz electrode. The results are represented by the mean ± standard error and the statistically significant differences (p=0.021) are indicated with (*). The athletes increases the P300 potential compared with non-athletes.

**Figure 4.**
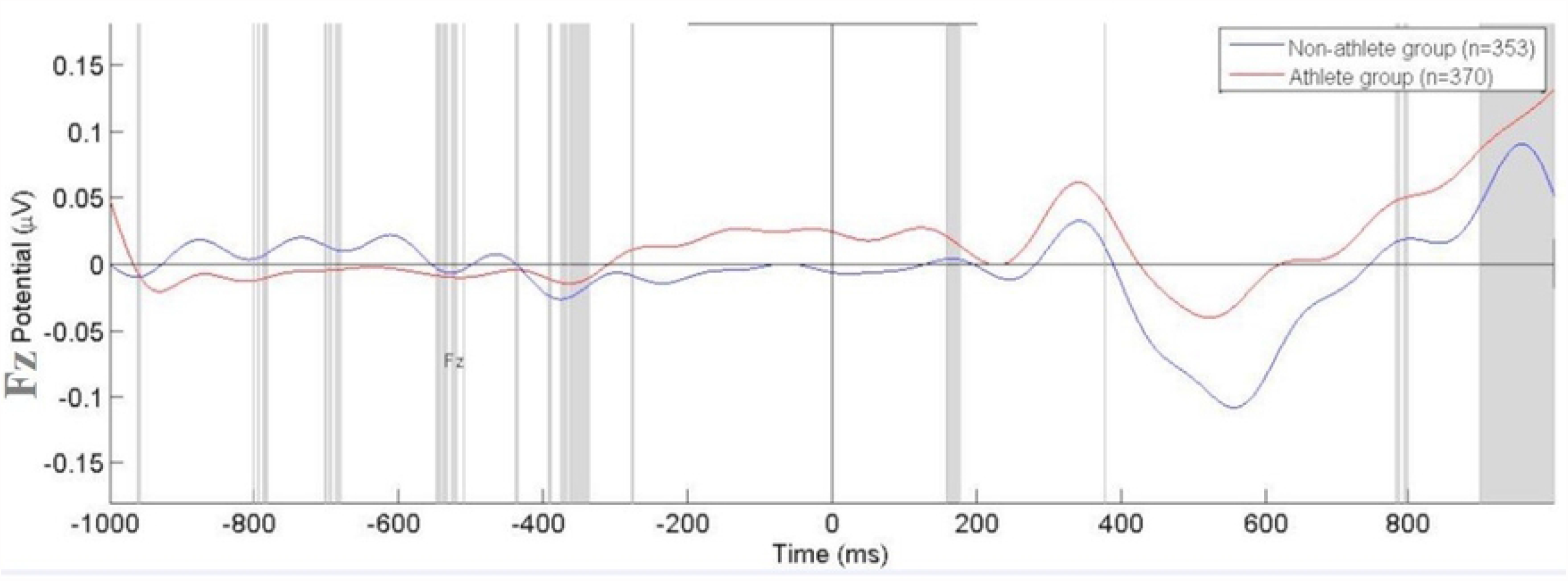
Mean Potential (μV) of visual P300 ERPs from the athletes group (red line) and non-athletes (blue line) in response to the Leeds stimulus obtained from the Fz electrode.

**Figure 5.**
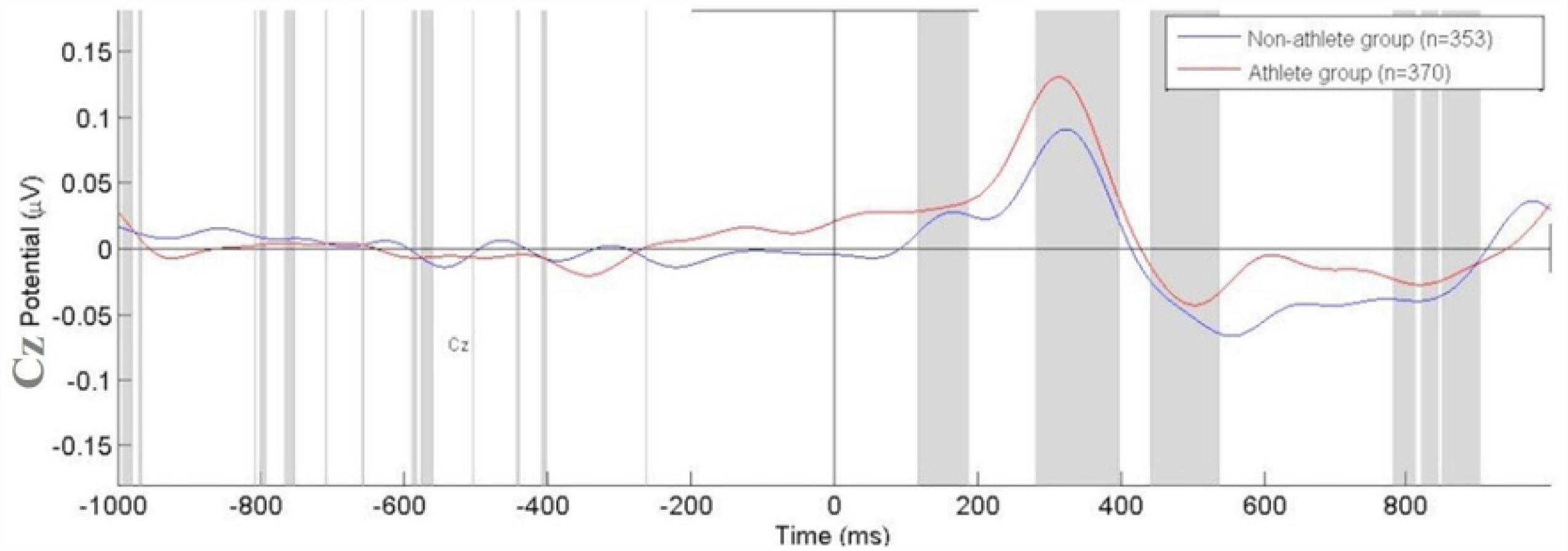
Mean Potential (μV) of visual P300 ERPs from the athletes group (red line) and non-athletes (blue line) in response to the Leeds stimulus obtained from the Cz electrode.

**Figure 6.**
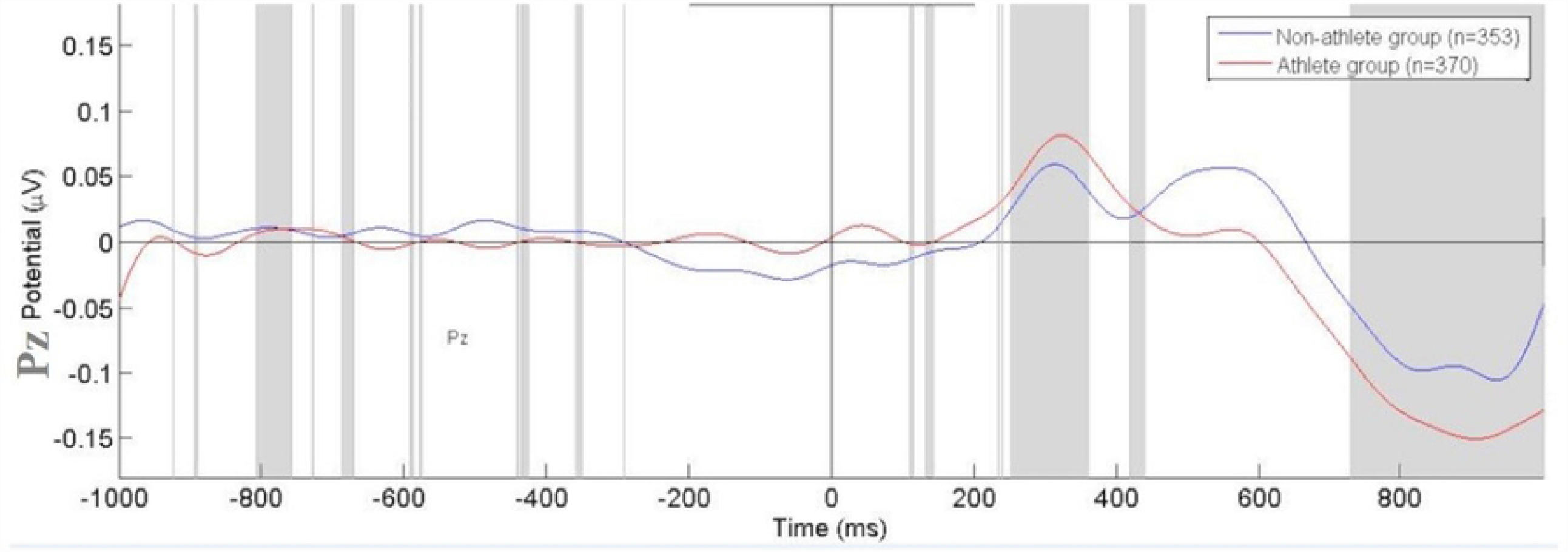
Mean Potential (μV) of visual P300 ERPs from the athletes group (red line) and non-athletes (blue line) in response to the Leeds stimulus obtained from the Pz electrode.

## 4. Discussion

We analyzed the differences in the electrocortical activity by P300 ERP component and reaction time comparing volleyball athletes and non-athletes. The hypothesis for differences in P300 was partial confirmed in the study. In relation reaction time, the findings showed a minimum effect, with a shorter reaction time for athletes.

The findings for reaction time corroborate with highly trained skill in high-level sports. We can consider that volleyball athletes need to process information from both central and peripheral vision, and the recurrence of training can adjust the visual focus improving reaction time during binocular, dominant eye and no dominant eye viewing conditions [26,34]. Since athletes respond faster to visual stimuli presented during a go/no go sensorimotor integration task. The recurrence of task training improve neural synchronization during the adjustment motor to respond to stimuli, whether visual, auditory or nociceptive.

The reaction in the decision-making triggered by visual stimuli is a part of dynamic interactions between personal experience and environment conditions and has been widely correlated with motor strategy [34]. Therefore, when reaction time is visualized in tasks of saccadic eye movement, there is a complex mechanism of inputs and outputs in the cortical regions [35,36].

Since reaction time functions involve recurrent interactions with external surroundings, the behavioral tasks can plausibly improve muscle response for a given objective. In this context, Giglia et al. [37] showed shorter reaction time in the open ability players group, which included volleyball athletes, than in the closed and sedentary skill group. This may be possible via the dopaminergic modulation in the connections of the basal ganglia with the cortical areas interconnectivity pattern of left and right hemispheres, which are related to the sensory-motor integration of the neurobiological process of attention [38]. Our inferences are supported on the ability of the saccadic eye movement tasks to encode changes in neuronal connections distributed in neural networks associated with adjustment and motor control (frontal, parietal and primary motor area). In addition, studies with badminton players have also shown a main effect of group, with the reaction time during a flanker task being lower for athletes than controls [38,39].

Previous studies show the effect of sports training in electroencephalographic results [3,4,38,39]. The hypothesis of differences in P300 ERPs between athletes and non-athletes during visual processing, states modulations in cortical energy consumption, since expert athletes can achieve better performance with less neural activity than non-athletes, which means neural efficiency [4].

The visual modulation might be the result of the specific requirements for a given sport training. Our findings demonstrated differences during the sensory processing only for the frontal midline, which may corroborate the precepts that athletes have more efficient neural response rates in the perception of visual stimuli [40,41]. The increase of the P300 amplitude in athletes than non-athletes suggest adjustment for the neurobiological functions of the attentional level for the stimulus, as well as the decision-making of response to the same [25,40,42].

This difference can evidence faster neural signal transmission in the visual focus [23]. The greater P300 potential during the saccadic eye movement paradigm tasks seems to occur due to lower cognitive demand to initiate the task, which is needed to create an internal model for planning, speed and execution [43]. This can be associated with working memory and attentional level [44], both related to the frontal area, which works as an integrating center of the neural inputs during visual perception task.

The visual architecture needs to provide information predicting when the stimuli will arrive [45]. In focus the volleyball, for effective response selection and action execution players must process and integrate a large amount of dynamic visual information, including flight information of the ball and kinetic information of the opponent. Thus, the P300 modulation in the frontal area might be the result of the specific requirements for a given sport training. In particular, the modulation of early sensory processing seems to be evident in athletes involved in ball sports requiring rapid responses to visual stimuli [3,21].

However, our findings no demonstrated differences and effects for motor and parietal interconnection area during rapid responses to visual stimuli. The findings of no difference in P300 activity can be evidenced by the difficulty of discriminating a target in a visual paradigm, could affect the scalp topography [18]. When targets stimuli occur in a series of more non-targets, a no significant P300 component is elicited over the parietal or central scalp areas [18,46].

The variability between genders, as well as heterogeneity within and between groups were limitations of this study. In addition, the sample size is a limiting factor, interfering with our results. Previous studies showed a low statistical power because the low signal-to-noise rates and the small sampling impacts the results [19,47]. However, the investigation of reaction time in volleyball athletes resulted in significant findings that may contribute to the knowledge of specific cognitive functions of this sport and may encourage the development of different training strategies to increase performance.

## 5. Conclusion

Our findings suggest that volleyball athletes demonstrate an allocation of attention to process the visual stimulus during the saccadic response task, and a shorter reaction time in the responses when compared to non-athletes. Therefore, this approach correlates the hypothesis of neural efficiency and the respective effects of sports training on the behavioral activity of athletes.

## References

[1] Sanchez-Lopez J, Fernandez T, Silva-Pereyra J, Martinez Mesa JA, Di Russo F. Differences in visuo-motor control in skilled vs. novice martial arts athletes during sustained and transient attention tasks: a motor-related cortical potential study. PLoS One. 2014 Mar 12;9(3):e91112.

[2] Sanchez-Lopez J, Silva-Pereyra J, Fernandez T. Sustained attention in skilled and novice martial arts athletes: a study of event-related potentials and current sources. PeerJ. 2016 Jan 26;4:e1614.

[3] Zwierko T, Lubiński W, Lesiakowski P, Steciuk H, Piasecki L, Krzepota J. Does athletic training in volleyball modulate the components of visual evoked potentials? A preliminary investigation. J Sports Sci. 2014;32(16):1519–28.

[4] Babiloni C, Marzano N, Infarinato F, Iacoboni M, Rizza G, Aschieri P, Cibelli G, Soricelli A, Eusebi F, Del Percio C. “Neural efficiency” of experts’ brain during judgment of actions: a high-resolution EEG study in elite and amateur karate athletes. Behav Brain Res. 2010 Mar 5;207(2):466–75.

[5] Abramov DM, Pontes M, Pontes AT, Mourao-Junior CA, Vieira J, Quero Cunha C, Tamborino T, Galhanone PR, deAzevedo LC, Lazarev VV. Visuospatial information processing load and the ratio between parietal cue and target P3 amplitudes in the Attentional Network Test. Neurosci Lett. 2017 Apr 24;647:91–96.

[6] Isoglu-Alkac U, Ermutlu MN, Eskikurt G, Yücesir İ, Demirel Temel S, Temel T. Dancers and fastball sports athletes have different spatial visual attention styles. Cogn Neurodyn. 2018 Apr;12(2):201–209.

[7] Del Percio C, Brancucci A, Vecchio F, Marzano N, Pirritano M, Meccariello E, Padoa S, Mascia A, Giallonardo AT, Aschieri P, Lino A, Palma E, Fiore A, Di Ciolo E, Babiloni C, Eusebi F. Visual event-related potentials in elite and amateur athletes. Brain Res Bull. 2007 Sep 14;74(1-3):104–12.

[8] Costanzo ME, VanMeter JW, Janelle CM, Braun A, Miller MW, Oldham J, Russell BA, Hatfield BD. Neural Efficiency in Expert Cognitive-Motor Performers During Affective Challenge. J Mot Behav. 2016 Nov-Dec;48(6):573–588.

[9] Liu T, Shao M, Yin D, Li Y, Yang N, Yin R, Leng Y, Jin H, Hong H. The effect of badminton training on the ability of same-domain action anticipation for adult novices: Evidence from behavior and ERPs. Neurosci Lett. 2017 Nov 1;660:6–11.

[10] Di Russo F, Pitzalis S, Spitoni G, Aprile T, Patria F, Spinelli D, Hillyard SA. Identification of the neural sources of the pattern-reversal VEP. Neuroimage. 2005 Feb 1;24(3):874–86.

[11] Zhang D, Ding H, Wang X, Qi C, Luo Y. Enhanced response inhibition in experienced fencers. Sci Rep. 2015 Nov 6;5:16282.

[12] Ludyga S, Mücke M, Andrä C, Gerber M, Pühse U. Neurophysiological correlates of interference control and response inhibition processes in children and adolescents engaging in open- and closed-skill sports. J Sport Health Sci. 2022 Mar;11(2):224–233.

[13] Helfrich RF, Knight RT. Cognitive neurophysiology: Event-related potentials. Handb Clin Neurol. 2019;160:543–558.

[14] Müller GR, Neuper C, Rupp R, Keinrath C, Gerner HJ, Pfurtscheller G. Event-related beta EEG changes during wrist movements induced by functional electrical stimulation of forearm muscles in man. Neurosci Lett. 2003 Apr 10;340(2):143–7.

[15] Müller-Putz GR, Scherer R, Pfurtscheller G, Rupp R. EEG-based neuroprosthesis control: a step towards clinical practice. Neurosci Lett. 2005 Jul 1-8;382(1-2):169–74.

[16] Neuper C, Pfurtscheller G. Post-movement synchronization of beta rhythms in the EEG over the cortical foot area in man. Neurosci Lett. 1996 Sep 20;216(1):17–20.

[17] Trejo LJ, Kubitz K, Rosipal R, Kochavi RL, Montgomery LD. EEG-Based Estimation and Classification of Mental Fatigue. Psychology. 2015; 6(6), 572–589

[18] Polich J, Comerchero MD. P3a from visual stimuli: typicality, task, and topography. Brain Topogr. 2003 Spring;15(3):141–52.

[19] Cavagnaro DR, Davis-Stober CP. A model-based test for treatment effects with probabilistic classifications. Psychol Methods. 2018 Dec;23(4):672–689.

[20] Bisagno E, Cadamuro A, Rubichi S, Robazza C, Vitali F. A developmental outlook on the role of cognition and emotions in youth volleyball and artistic gymnastics. Front Psychol. 2022 Aug 10;13:954820.

[21] Zwierko T, Lubiński W, Lubkowska A, Niechwiej-Szwedo E, Czepita D. The effect of progressively increased physical efforts on visual evoked potentials in volleyball players and non-athletes. J Sports Sci. 2011 Nov;29(14):1563–72.

[22] Rizzolatti G, Riggio L, Dascola I, Umiltá C. Reorienting attention across the horizontal and vertical meridians: evidence in favor of a premotor theory of attention. Neuropsychologia. 1987;25(1A):31–40.

[23] Ozmerdivenli R, Bulut S, Bayar H, Karacabey K, Ciloglu F, Peker I, Tan U. Effects of exercise on visual evoked potentials. Int J Neurosci. 2005 Jul;115(7):1043–50. doi: 10.1080/00207450590898481. PMID: 16051549.

[24] Fisher P, Schenk T. Temporal order judgments and presaccadic shifts of attention: What can prior entry teach us about the premotor theory? J Vis. 2022 Nov 1;22(12):6.

[25] Sanfim A, Velasques B, Machado S, Arias-Carrión O, Paes F, Teixeira S, Santos JL, Bittencourt J, Basile LF, Cagy M, Piedade R, Sack AT, Nardi AE, Ribeiro P. Analysis of slow- and fast-α band asymmetry during performance of a saccadic eye movement task: dissociation between memory- and attention-driven systems. J Neurol Sci. 2012 Jan 15;312(1-2):62–7.

[26] Bittencourt J, Velasques B, Teixeira S, Basile LF, Salles JI, Nardi AE, Budde H, Cagy M, Piedade R, Ribeiro P. Saccadic eye movement applications for psychiatric disorders. Neuropsychiatr Dis Treat. 2013;9:1393–409.

[27] Bowling AC, Lindsay P, Smith BG, Storok K. Saccadic eye movements as indicators of cognitive function in older adults. Neuropsychol Dev Cogn B Aging Neuropsychol Cogn. 2015;22(2):201–19.

[28] Oldfield RC. The assessment and analysis of handedness: the Edinburgh inventory. Neuropsychologia. 1971 Mar;9(1):97–113.

[29] Marin Rueda FJ, Noronha APP, Sisto FF, Bartholomeu D. Evidência de validade de construto para o teste de atenção sustentada. Psicologia: Ciência e Profissão. 2008; 28(3), 494–505. https://doi.org/10.1590/S1414-98932008000300005

[30] Szurhaj W, Derambure P, Labyt E, Cassim F, Bourriez JL, Isnard J, Guieu JD, Mauguière F. Basic mechanisms of central rhythms reactivity to preparation and execution of a voluntary movement: a stereoelectroencephalographic study. Clin Neurophysiol. 2003 Jan;114(1):107–19.

[31] Gongora M, Peressuti C, Velasques B, Bittencourt J, Teixeira S, Arias-Carrión O, Cagy M, Ribeiro P. Absolute Theta Power in the Frontal Cortex During a Visuomotor Task: The Effect of Bromazepam on Attention. Clin EEG Neurosci. 2015 Oct;46(4):292–8.

[32] Ferguson C. An effect size primer: A guide for clinicians and researchers. Professional Psychology: Research and Practice. 2009; 40(5), 532–538.

[33] Jascaniene N, Nowak R, Kostrzewa-Nowak D, Kolbowicz M. Selected aspects of statistical analyses in sport with the use of STATISTICA software. Central European Journal Sport Sciences Medica. 2013; 3, 3–11.

[34] Avanzino L, Pelosin E, Vicario CM, Lagravinese G, Abbruzzese G, Martino D. Time Processing and Motor Control in Movement Disorders. Front Hum Neurosci. 2016 Dec 12;10:631.

[35] Fontes R, Ribeiro J, Gupta DS, Machado D, Lopes-Júnior F, Magalhães F, Bastos VH, Rocha K, Marinho V, Lima G, Velasques B, Ribeiro P, Orsini M, Pessoa B, Leite MA, Teixeira S. Time Perception Mechanisms at Central Nervous System. Neurol Int 2016; 8(1):5939).

[36] Marinho V, Oliveira T, Rocha K, Ribeiro J, Magalhães F, Bento T, Pinto GR, Velasques B, Ribeiro P, Di Giorgio L, Orsini M, Gupta DS, Bittencourt J, Bastos VH, Teixeira S. The dopaminergic system dynamic in the time perception: a review of the evidence. Int J Neurosci. 2018 Mar;128(3):262–282.

[37] Giglia G, Brighina F, Zangla D, Bianco A, Chiavetta E, Palma A, Fierro B. Visuospatial attention lateralization in volleyball players and in rowers. Percept Mot Skills. 2011 Jun;112(3):915–25.

[38] Wang CH, Tu KC. Neural Correlates of Expert Behavior During a Domain-Specific Attentional Cueing Task in Badminton Players. J Sport Exerc Psychol. 2017 Jun 1;39(3):209–221.

[39] Del Percio C, Iacoboni M, Lizio R, Marzano N, Infarinato F, Vecchio F, Bertollo M, Robazza C, Comani S, Limatola C, Babiloni C. Functional coupling of parietal α rhythms is enhanced in athletes before visuomotor performance: a coherence electroencephalographic study. Neuroscience. 2011 Feb 23;175:198–211.

[40] Kamijo K, Takeda Y, Takai Y, Haramura M. Greater aerobic fitness is associated with more efficient inhibition of task-irrelevant information in preadolescent children. Biol Psychol. 2015 Sep;110:68–74.

[41] Pontifex MB, Raine LB, Johnson CR, Chaddock L, Voss MW, Cohen NJ, Kramer AF, Hillman CH. Cardiorespiratory fitness and the flexible modulation of cognitive control in preadolescent children. J Cogn Neurosci. 2011 Jun;23(6):1332–45.

[42] Retzlaff PD, Morris GL. Event-related potentials during the Continuous Visual Memory Test. J Clin Psychol. 1996 Jan;52(1):43–7.

[43] Levy BJ, Wagner AD. Cognitive control and right ventrolateral prefrontal cortex: reflexive reorienting, motor inhibition, and action updating. Ann N Y Acad Sci. 2011 Apr;1224:40–62.

[44] Melo HM, Nascimento LM, Mello VO, Takase E. Alpha (8-12Hz) influence on reaction time in inhibitory control task. Neuropsychology Latinoamericana. 2017; 9(2):33–43.

[45] Le Runigo C, Benguigui N, Bardy BG. Visuo-motor delay, information-movement coupling, and expertise in ball sports. J Sports Sci. 2010 Feb;28(3):327–37.

[46] Posner MI, Petersen SE. The attention system of the human brain. Annu Rev Neurosci. 1990;13:25–42.

[47] Dowding I, Haufe S. Powerful Statistical Inference for Nested Data Using Sufficient Summary Statistics. Front Hum Neurosci. 2018 Mar 19;12:103.

